# Complete loss of CASK causes severe ataxia through cerebellar degeneration in human and mouse

**DOI:** 10.1101/2021.03.22.436280

**Authors:** Paras A Patel, Julia Hegert, Ingrid Cristian, Alicia Kerr, Leslie EW LaConte, Michael A Fox, Sarika Srivastava, Konark Mukherjee

**Affiliations:** Translational Biology, Medicine, and Health Graduate Program, Fralin Biomedical Research Institute, Roanoke, VA 24016, USA; Center for Neurobiology Research, Fralin Biomedical Research Institute, Roanoke, VA 24016, USA; Virginia Tech, Department of Biological Sciences, Blacksburg, VA 24060, USA; Virginia Tech, School of Neuroscience, Blacksburg, VA, 24060, USA; Orlando Health, APH, MP 331, USA; Virginia Tech Carilion School of Medicine, Department of Basic Science Education. Roanoke, VA 24016, USA; Department of Internal Medicine, Virginia Tech Carilion School of Medicine. Roanoke, VA 24016, USA; Department of Psychiatry, Virginia Tech Carilion School of Medicine, Roanoke, VA 24016, USA

## Abstract

Heterozygous loss of X-linked genes like CASK and MeCP2 (Rett syndrome) causes neurodevelopmental disorders (NDD) in girls, while in boys such loss leads to profound encephalopathy. The cellular basis for these disorders remains unknown. CASK is presumed to work through the Tbr1-reelin pathway in neuronal migration during brain development. Here we report our clinical and histopathological analysis of a deceased 2-month-old boy with a CASK-null mutation. We demonstrate that although smaller in size, the CASK-null human brain exhibits normal lamination without defective neuronal differentiation, migration, or axonal guidance, excluding the role of reelin in CASK-linked pathology. The disproportionately hypoplastic cerebellum in humans without CASK expression is associated with cerebellar astrogliosis, a marker for neuronal loss. Cerebellum-specific deletion in mouse confirms a post-developmental degeneration of cerebellar granular neurons that results in a small cerebellum. Mechanistically, cerebellar hypoplasia in CASK mutation thus results from neurodegeneration rather that developmental defects. Zygosity-pathology correlation suggests that NDDs like CASK mutation and Rett syndrome are pathologically neurodegenerative; however, random X-chromosome inactivation in the typical heterozygous mutant girls results in 50% of cells expressing the functional gene, resulting in a non-progressive pathology, whereas complete loss of the only allele in boys leads to unconstrained degeneration and encephalopathy.

**One sentence summary of study:** CASK loss causes cerebellar degeneration. The authors have declared that no conflict of interest exists.

## Introduction

Heterozygous mutations in certain X-linked genes (e.g., CDKL5, MeCP2 in Rett syndrome, and CASK in MICPCH (<u>m</u>ental retardation and m<u>ic</u>rocephaly with <u>p</u>ontine and <u>c</u>erebellar <u>h</u>ypoplasia (OMIM: 300749)) are linked to postnatal microcephaly in girls ^1^. Hemizygous mutations in these same genes give rise to progressive epileptic encephalopathy and lethality in boys ^2–4^. Rett syndrome was the first such disorder to be reported; it was described as a cerebral atrophic syndrome by Andreas Rett in 1966 ^5^. Until the 1990s, Rett syndrome was considered a neurodegenerative disorder ^6^. With the discovery, however, of the MeCP2 gene association, postmortem autopsy observations, and the development of preclinical models, focus shifted to dendritic morphology and synapse development and dysfunction, resulting in the re-classification of Rett syndrome as a neurodevelopmental disorder ^7 8^. Studies on the cellular pathology associated with MeCP2 loss in boys with epileptic encephalopathies have, however, been limited ^9^.

MICPCH is also considered to be a neurodevelopmental disorder that occurs due to heterozygous mutations in the X-linked gene CASK (calcium/calmodulin-dependent serine protein kinase) in girls. Despite the microcephaly associated with CASK mutation being described as postnatal and progressive, females with MICPCH grow into adulthood, often with an intellectual disability that is non-progressive ^10–14^. Such mutations in hemizygous males are, however, lethal. These boys exhibit epileptic encephalopathy with pronounced cerebellar hypoplasia and progressive supratentorial atrophy ^4, 15, 16^. Regression of motor skills has also been noted in a girl with MICPCH in adolescence ^17^. The cellular pathology of CASK-linked disorders remains uncertain. This problem is exacerbated by the fact that CASK-null mice die within hours of birth and do not exhibit a difference in brain size or morphology from their wild type littermates at birth ^18^. Based on the standard Theiler developmental staging of mice ^19^, the immediate postnatal period of mice best parallels the third trimester of human embryonic development (Carnegie staging; ^20^), making any interpretation of postnatal brain pathology difficult.

Although often considered to be a component of presynaptic terminals, CASK in fact is ubiquitously expressed in the body and has been implicated in a variety of functions ^21, 22^. Outside of the brain, CASK has been shown to participate in cell proliferation ^23^, cell polarization ^24^, gap junctions and wound healing ^25^, insulin secretion and signaling ^26, 27^, hypoxia response ^28, 29^, renal development and disease ^30, 31^, spermatogenesis and sperm motility ^32, 33^, and cardiac conductivity ^34, 35^, to name a few examples. In the brain, CASK has been examined as both a pre- and post-synaptic molecule ^36, 37^. It has also been suggested that CASK is involved in protein trafficking via its interaction with SAP97 ^38, 39^. CASK is proposed to be involved in axonal branching ^40^, dendritic arborization ^41^, dendrite spinogenesis ^42^ and synaptogenesis ^43^. Thus, many hypotheses as to why loss of CASK leads to defects in brain development can be proposed.

CASK also has a function in regulating gene transcription ^44–46^. It has been suggested that CASK translocates to the nucleus, where it regulates the function of T-box transcription factor (Tbr-1) ^46, 47^. It is proposed that CASK forms a ternary complex together with CINAP (CASK-interacting nucleosome assembly protein) and Tbr-1 to induce expression of molecules such as reelin that play a crucial role in brain development ^45^. Reelin is a secreted extracellular molecule critical for neuronal migration^48^. Indeed, both in the reeler mice and Tbr-1 knockout mice, defects in proper lamination of cortex are seen ^49, 50^. In addition to the cortex, reeler mice also display a hypoplastic disorganized cerebellum with defects in neuronal migration and suppressed neurogenesis ^49^. The neurodevelopmental function of CASK has been specifically attributed to the CASK’s interaction with Tbr-1and presumed regulation of reelin expression ^13, 51, 52^.

Here, we report a detailed clinical description and autopsy findings from a 2-month-old boy harboring the CASK null mutation R27*. Although the brain is small, the clear presence of tertiary gyri that form near term, as well as proper cortical and cerebellar lamination, argue against defects in neuronal migration; instead we uncover evidence of neurodegeneration suggested by reactive astrogliosis in the cerebellum. We then design and execute a genetic experiment in mice that provides conclusive evidence that loss of CASK indeed produces neurodegeneration in the cerebellum. Most pontocerebellar hypoplasias (PCH) are progressive, but based on the postnatal brain growth pattern, it has been hypothesized that MICPCH has a distinct pathogenic mechanism^53^. We instead provide evidence that mechanistically, MICPCH in girls with heterozygous CASK mutations is also degenerative, and the non-progressive course of MICPCH is dictated by uniqueness of the X-linked inheritance pattern in which 50% of brain cells express the normal gene.

## Results

### Complete CASK loss in humans causes profound neurodevastation and cerebellar atrophy

MICPCH subjects with heterozygous CASK mutations are known to live past their 30s. *Cask*^+/-^ female mice are fertile beyond 6 months. We have allowed four *Cask*^+/-^ mice to age more than two years, considered to be old for mice. All four mice survived to that age without adverse events. We did not observe any obvious phenotypes in these aged mice compared to wild-type littermates. The cerebellum displayed the typical layers and configuration without severe deterioration, indicating that the disorder is non-progressive (Supplemental Figure 1).

Null mutation of *CASK* in mice is, however, lethal and in boys, produces progressive encephalopathy. Due to the early lethality of *Cask* null mice, the postnatal pathology of complete *Cask* loss has been difficult to study to date. Here we describe detailed clinical findings and autopsy results from a 2-month-old boy with a *CASK* null mutation who expired due to hypoventilation and neurogenic respiratory failure. A copy number variation study was unremarkable, but next generation sequencing of genes revealed a c.79C>T (p.Arginine27Ter) *CASK* mutation in exon 2 (Figure 1A). This *CASK* mutation introduces a stop codon in the very N-terminus of the CASK protein, precluding expression of any splice variant of CASK (Supplemental Figure 3). Magnetic resonance imaging (MRI) indicated normal lateral and third ventricles with an elongated fourth ventricle. The cerebellum appeared markedly hypoplastic without a vermis. The small posterior fossa was filled with fluid (Figure 1B). The corpus callosum was thin but present without any midline shift, and myelination was delayed for age. The cavum septum pellucidum seemed to be more prominent. No heterotopic cells were noted in any area, but there was some degree of smoothening, particularly of the frontal cortex. The brain stem appeared to be extremely thin.

**Figure 1.**
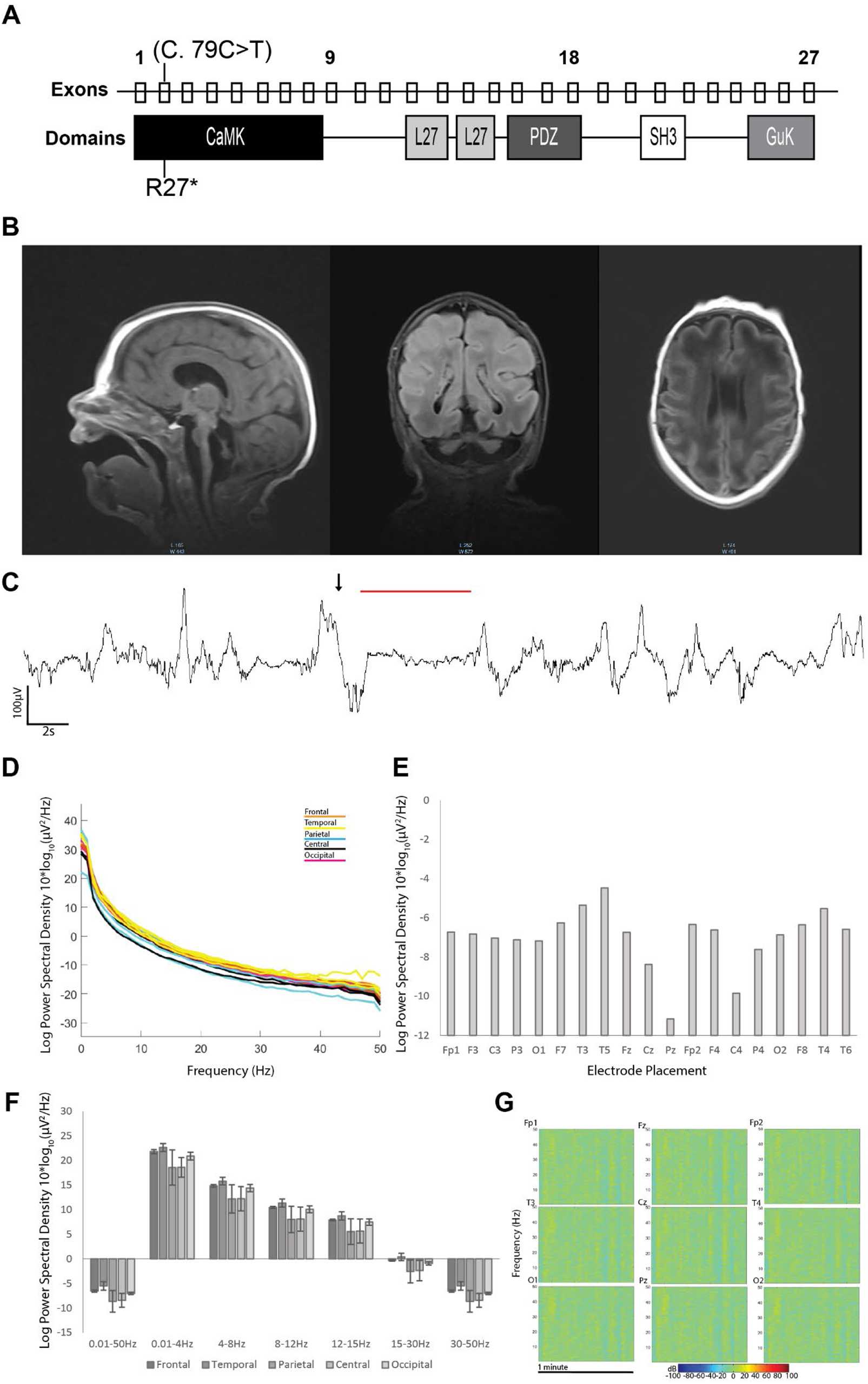
The R27Ter mutation results in early truncation of *CASK,* pontocerebellar hypoplasia and slowed EEG in a human subject. A) Location of the R27Ter mutation in the male decedent; top row: exons of the human *CASK* gene; bottom row: corresponding CASK protein domains. B) Brain MRI scan obtained at 2 weeks revealed severely diminished cerebellar and pontine size with otherwise normal formation of cortical gyri; sagittal, coronal, and transverse planes can be seen from left to right, respectively. C) A representative example of discontinuous EEG pattern (arrow and red line in the EEG trace); more examples of burst suppression can be found in Supplemental Figure 2. D) Power spectral density curves in the 0.01-50Hz range for each electrode independently; colors correspond to underlying cortical location of a given electrode. E) Quantification of mean power spectral density for each electrode in the entire 0.01-50Hz range. F) Mean power spectral density divided into biologically relevant frequency bands of delta, theta, alpha, beta, and low gamma divided by lobe; error bars represent standard deviation between electrodes within a given lobe. G) Representative time frequency plots of 1 minute of the recording by electrode position.

Video electroencephalographic (vEEG) monitoring was done both during awake and sleeping states. Awake-state background EEG displayed a burst-suppression pattern with variable amounts of bursts and suppressions (Figure 1C and Supplemental Figure 2). This EEG pattern is typical of Ohtahara syndrome, a devastating epileptic encephalopathy, that usually co-occurs with *CASK*-null mutations ^4, 15, 16^. The burst phase was dominated by a mixture of theta and delta waves. Overall, the EEG retained its symmetry in both hemispheres but was discontinuous. No electroclinical seizures were observed during the period of recording, although intermittent and independent sharp waves were observed, predominantly in the right temporal and occipital region. The sleep EEG was similar to the waking EEG and included burst-suppression signals. A spectral analysis of the entire epoch revealed skewing towards lower frequency with delta and alpha power dominating the spectra (Figure 1D-G).

At autopsy, head circumference was 32.7 cm, with a 37.0 cm crown-rump length and crown-heel length of 51.0 cm. The decedent was small for his age, and the brain weight was 300.8 grams, which is 60% of what is expected at this age (Figure 2A). Except for lung, heart, and spleen, most other organs were smaller than expected but had an overall normal gross appearance (Figure 2A, Supplemental Figure 5). The brain was well formed with normal gyri formations in the cerebral hemispheres. Tertiary gyri were present, and there was no evidence of polymicrogyria or other abnormal configuration. The Sylvian fissure was well formed, and the leptomeninges were clear (Figure 2B). Vascularization, including the circle of Willis, was normally formed. The central part of the cerebral hemispheres was edematous, and the septum cavum pellucidum was present (0.9 cm in vertical length). The basal ganglia displayed a normal architecture bilaterally. The left hippocampus was also architecturally normal with a serpiginous appearance. The right hippocampus had a blurred appearance (Supplemental Figure 4). The thalamus was normally formed and firm. The lateral ventricles were not dilated; the midbrain was very small with a patent but pinpoint cerebral aqueduct, and the fourth ventricle was slit-like. The cerebellum and the pons were markedly hypoplastic (Figure 2C, Supplemental Figure 4). The cerebellum, despite hypoplasia, had a normal configuration but did not exhibit the usual folia. There was no evidence for heterotopia of cells. The anterior vermis was not identifiable and appeared to be membrane-like; cerebellar hemispheres were thin, flattened and firm. The spinal cord was of uniform caliber and had no obvious pathology.

**Figure 2.**
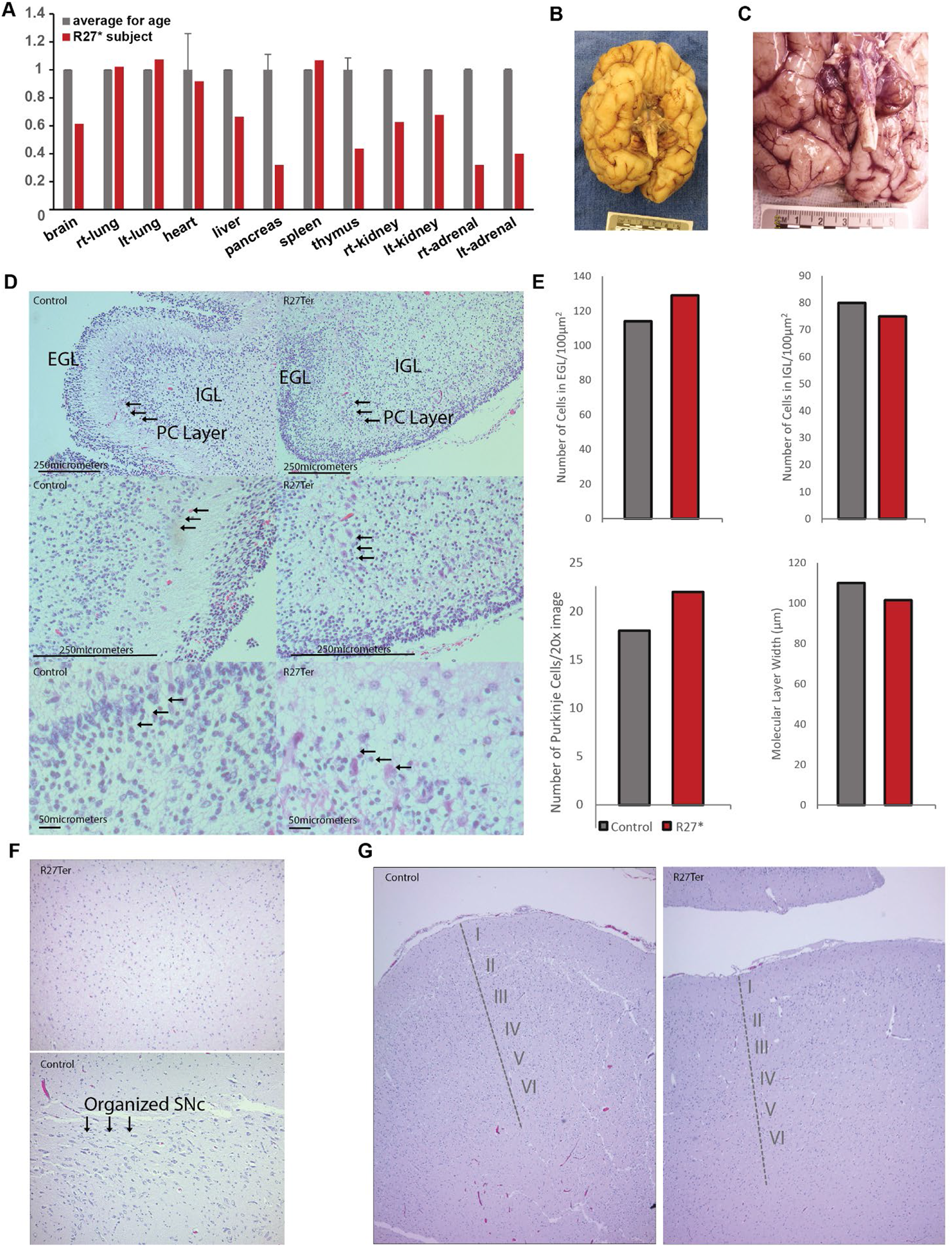
Absence of CASK leads to smaller organs including brain with normal histology of cerebellum and no defect in lamination or cellular migration. A) Organ weight of the decedent relative to average weight for the age ^79^. B) Gross image of the underside of the brain of the decedent showing normal cortical gyri development with a severely diminished cerebellar volume. C) Close-up of image in (B). D) H&E staining of cerebellar cortex of the R27Ter subject and of a 46-day-old infant female (died of unrelated cause), indicating proper lamination of cerebellar cortex with external granular layer (EGL), internal granular layer (IGL), and Purkinje cell layers; arrows indicate properly aligned Purkinje cells. E) Quantification of granule cells in the internal and external granular layers, Purkinje cells, and width of the molecular layer between the R27Ter subject and a 20-day-younger female who died of an unrelated cause. F) H&E-stained substantia nigra of the decedent and control showing lack of an organized substantia nigra in the absence of CASK. Arrows in the control brain indicate neuronal cells in the substantia nigra (purple color). G) H&E-stained cortex of R27Ter subject and control demonstrating proper cortical lamination in the presence and absence of CASK.

### Absence of CASK does not affect neuronal migration, axonal guidance, or lamination in humans but may promote neuronal loss

Histologically, the cerebellum itself displayed proper cellular organization, with a defined external granular layer (EGL), molecular layer, and internal granular layer (IGL). There was a uniform single layer of Purkinje cells between the molecular layer and the internal granular layer (Figure 2D, E). A proper migratory pattern of granule cells was visible and appropriate for age. The white matter was poorly organized, and the dentate nucleus was absent. The midbrain consisted of astrocytic cells with pink cytoplasm and some neuronal cells, however no organized substantia nigra was noted (Figure 2F). Sections of the cortex indicated orderly and proper neuronal migration; the germinal matrix was appropriately thinned for this age. The white matter tracts were discreet and adequate for this age (Supplemental Figure 4). The basal ganglia displayed normal numbers of neurons. The hippocampi were properly organized with uniform neuronal populations in all CA (cornu ammonis) zones. The midbrain and pons displayed corticospinal tracts. The cerebral aqueduct was patent and dilated. Within the pons, the pontine decussation was seen and the locus coeruleus properly formed. The medullary olives were poorly formed, and the fourth ventricle was widely patent. The spinal cord was unremarkable with adequate anterior horn cells and uniform radiating column. The central canal was patent throughout (Supplemental Figure 4).

Histologically, almost all organs including the bone marrow, heart, and intestines were unremarkable. The endocrine glands also appeared normal, except for the adrenal cortex, which was thinned out. The kidneys had appropriate and orderly glomerular and tubular development (Supplemental Figure 5C, D). The heart rate varied between 90 and 150 beats per minute and displayed a sinus rhythm (Supplemental Figure 6).

Data clearly indicate that although CASK loss affects the size of the brain globally, the cerebellum and brainstem are disproportionately affected. Both in murine models and the human subject, early lethality is likely linked with the dysfunction of the brainstem leading to respiratory failure. *Cask* null mice display hypoventilation and die within hours of birth although the brain is of normal size and properly laminated at death ^18^.

The normal lamination and configuration of the brain in both CASK null humans and mice suggests that the histological pathology related to CASK loss is likely to be neurodegenerative, with neuronal loss. One of the most common hallmarks of neuronal damage and neuronal loss is reactive gliosis. We therefore next evaluated the cerebellum of the decedent for the ability to form synapses and for evidence of astrogliosis (Figure 3). Previous studies in the murine model have shown that CASK loss-of-function does not negatively impact synapse formation ^18, 54^, and in the human cerebellum evaluated here, immunostaining revealed that levels of the synaptic marker synaptophysin in the decedent’s cerebellum were similar to levels observed in an age-matched control cerebellum (Figure 3A, B). GFAP staining of the cerebellum to detect astrogliosis, however, indicates that, compared to the control, the decedent exhibits ∼5-fold higher amounts of GFAP immunoreactivity, specifically in the IGL (Figure 3A, C, D). In fact, large reactive astrocytes in the IGL were readily observed (Figure 3C). Together these data suggest that loss of CASK produces delayed neurodegenerative changes, causing the CASK-linked phenotype to typically manifest postnatally. We finally investigated myelination within the cerebellar cortex using FluoroMyelin lipid staining (Figure 3E) and observed that the myelination pattern exhibited disturbed arrangement within the IGL of the R27Ter subject, with discrete myelinated tracts, in contrast to the diffuse mesh-like myelin staining observed in the control subject. The histological features thus indicate that the disorganized white matter described earlier may be secondary to ongoing cerebellar grey matter degeneration. Thus our observations support the notion that loss of CASK induces cerebellar cortical degeneration, specifically in the IGL. To test this idea, we next employed murine genetic experiments, where CASK is deleted in a temporally and spatially specified manner using Cre-LoxP-mediated gene excision.

**Figure 3.**
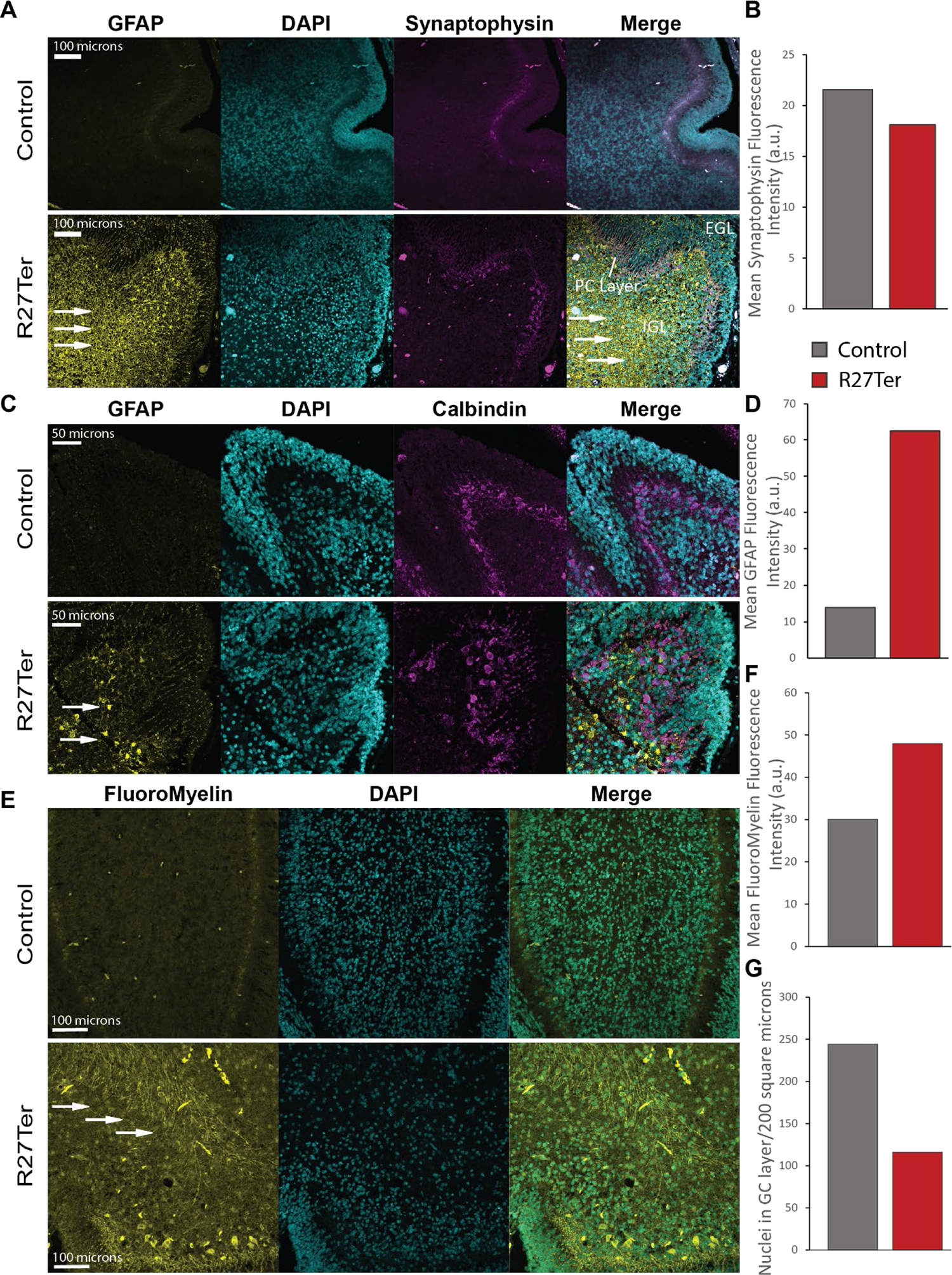
The R27Ter mutation is associated with pronounced astrogliosis in cerebellum, and defects in white matter. (A) Representative images of GFAP and synaptophysin immunostaining in the R27Ter and age-matched control human cerebellum; from left to right: GFAP, DAPI, synaptophysin, merge. Arrows indicate increased GFAP immunoreactivity. (B) Quantification of synaptophysin counterstain, demonstrating similar staining intensity in control and R27Ter subject. (C) Representative images of GFAP and calbindin immunostaining in the R27Ter and age-matched control human cerebellum; from left to right: GFAP, DAPI, calbindin, merge. Arrows indicate increased GFAP immunoreactivity. (D) Quantification of GFAP immunostaining fluorescence intensity between R27Ter cerebellum and control cerebellum. (E) Fluorescent labelling of myelin using FluoroMyelin indicating frayed white matter with individually visible axons in the R27Ter subject compared to the diffuse staining observed in the age-matched control; from left to right: FluoroMyelin staining, DAPI, merge. (F) Quantification of FluoroMyelin fluorescence intensity. (G) Quantification of DAPI+ nuclei in the granular layer indicating a decreased number of cells.

### *Calb2*-Cre targets post-migratory granule cells and a subset of Purkinje cells in the cerebellum

Previous neuroimaging data and the comprehensive CASK null brain histological autopsy results presented here clearly indicate that within the brain, loss of CASK is likely to disproportionately affect the hindbrain including the brainstem and cerebellum. In particular, CASK-linked lethality most likely results from effects on the brain stem. Our focus, therefore, in the study presented here is to evaluate the long-term effect of CASK loss in the cerebellum. We have previously demonstrated that CASK loss likely does not affect cerebellar development. We reached this conclusion by examining three different mouse constructs: 1) <u>pan-neuronal *Cask* knockout mice,</u> which die before P24 (postnatal day 24) but exhibit normal cerebellar formation and lamination ^54^; 2) <u>Purkinje cell-specific knockout mice,</u> which display normal development and motor function ^54^; and finally, 3) <u>mice with *Cask* deletion in a distributed subpopulation of granule cells,</u> which do not exhibit altered cell migration or survival ^54^. There are two critical reasons that conclusions about cerebellar development from these previous experiments must be tempered: 1) for each of these mouse types, *Cask* was deleted only in small subset of cells in the cerebellum ^55–57^; and 2) we did not study the long-term effect of *Cask* deletion in the cerebellum. To address these gaps and examine the role of CASK in the cerebellum over longer time periods, we have devised a method to delete CASK from most cerebellar cells in a manner that does not produce lethality in mice. To do so, we chose a mouse line in which Cre-recombinase is driven by an endogenous promoter of the signaling molecule *Calb2* (calretinin/calbindin2), reported earlier ^58^. The choice of *Calb2*-Cre was made instead of a promoter such as *Math1*, a transcription factor, because *Math1* turns on earlier and is also expressed in the brain stem, which could contribute to lethality. It has been shown that *Calb2* expresses in nearly all granule cells in the cerebellum ^59, 60^, but the exact timing of initiation of *Calb2*-Cre gene recombination in granule cells was not known. There have also been conflicting reports about the expression of *Calb2* in the Purkinje cells within the cerebellum ^59, 60^. We therefore first tested the recombination specificity of *Calb2*-Cre in mice at ages when the cerebellum is still developing and displays both the EGL and IGL (P8 and P15). We crossed *Calb2*-Cre mice with Cre-recombination indicator mice (LSL-tdTomato) (Figure 4A). The distribution of the tdTomato-expressing neurons serves as a proxy for CASK deletion when *Calb2*-Cre mice are crossed with *Cask*^floxed^ mice in parallel (Figure 4B). Our data indicate that *Calb2*-Cre is active in the cerebellum as early as P8. By P8, recombination was observed in granule cells but only after migration into the IGL; recombination was also observed in many Purkinje cells (Figure 4C). By P15, Calb2-Cre already exhibited robust recombination in many parts of the brain and in the entirety of post-migratory granule cells. Dense cellular distribution with recombination was seen in the cerebellum, hippocampus, striatum and olfactory bulb. Sparsely distributed cells were observed throughout the brain, including the cortex (Figure 4D, E). The brainstem displayed minimal recombination, with sparsely tdTomato-labeled cells. Within the cerebellum, all granular cells in the IGL and a subset of Purkinje cells were positive for recombination at P15. Cells in the EGL, however, did not display any recombination, indicating that *Calb2-*Cre-driven recombination occurs only after migration of granule cells (Figure 4E). Our data thus indicate that *Calb2*-Cre specifically leads to deletion of CASK both in a subset of Purkinje cells and in granule cells within the IGL by P15 and is not likely to affect the brainstem or its function.

**Figure 4.**
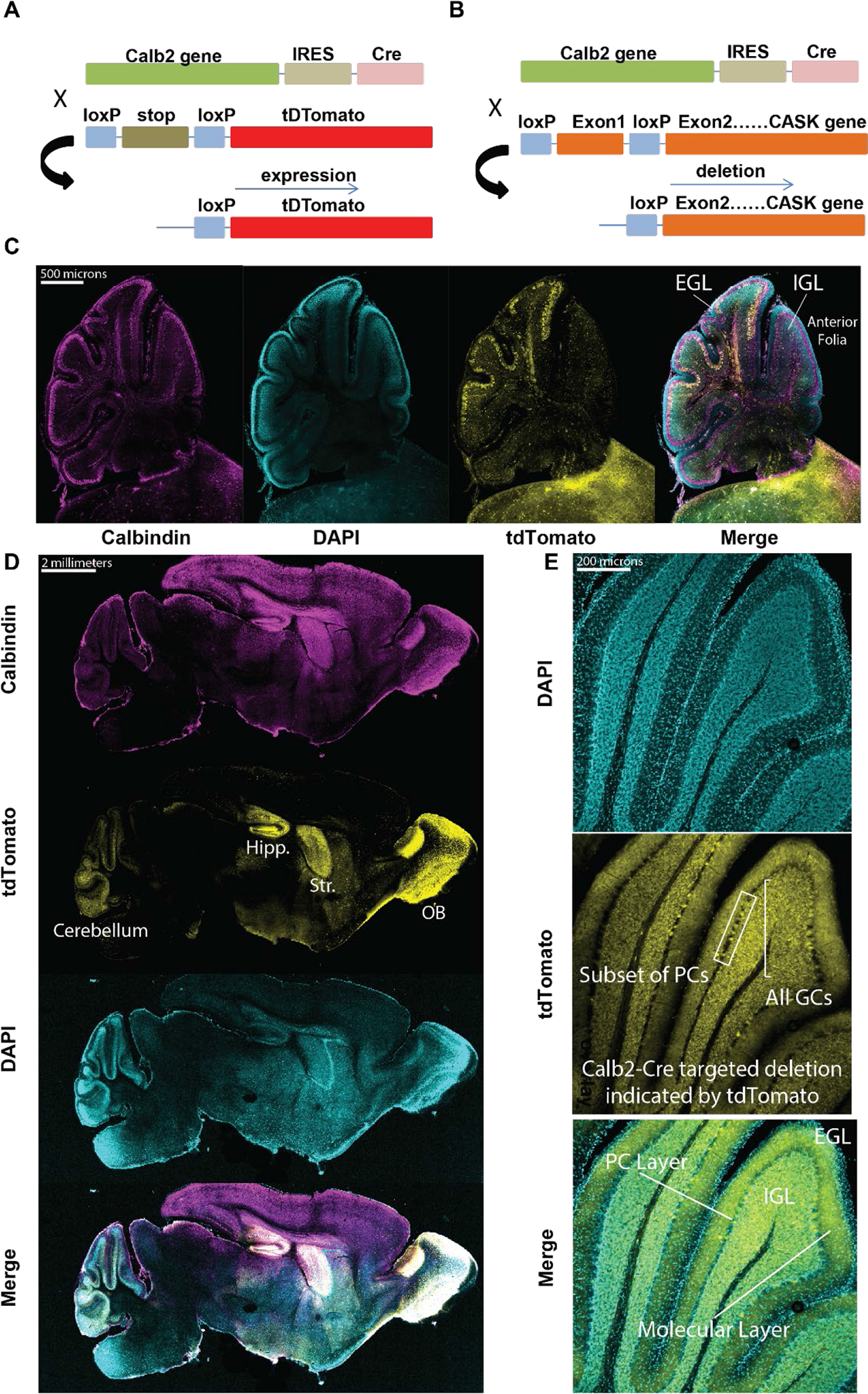
*Calb2*-Cre expresses only in post-migratory cerebellar granule cells and a subset of Purkinje cells in the anterior folia. (A) Generation of LoxP-STOP-LoxP tdTomato reporter mice for determination of cell-type and age-specific Cre-mediated recombination; tdTomato expression serves as a proxy for CASK deletion in subsequent panels. (B) Schematic of breeding strategy used to selectively delete *Cask* from cerebellar cells. (C) Sagittal section of cerebellum at P8, demonstrating recombination in a subset of Purkinje cells and cells in the internal granular layer, but not in granule cells of the external granular layer; from left to right: tdTomato, DAPI, merge. (D) 20x images of a sagittal section of whole mouse brain at P15, demonstrating recombination in several brain regions, notably the cerebellum, olfactory bulb (OB), hippocampus (Hipp.), and striatum (Str.) after development; from top to bottom: calbindin, tdTomato, DAPI, merge. (E) Higher magnification images of cerebellar folia at P15, demonstrating recombination in virtually all cells in the granular layer but only in a small subset of Purkinje cells; box indicates a sample of the subpopulation of Purkinje cells expressing tdTomato reporter while the bracket indicates all granule cells observed expressing tdTomato. The Purkinje cell (PC) layer, molecular layer, IGL, and EGL are indicated for one folium for anatomical orientation.

### Deletion of CASK from cerebellar neurons results in later-onset progressive degeneration of the cerebellum and severe ataxia

We next examined mice from crosses of the *Calb2*-Cre and *Cask*^floxed^ lines. It has been shown previously that the *Cask*^floxed^ mouse is a hypomorph that expresses ∼40% CASK, likely due to a phenomenon known as selection cassette interference ^18^. *Cask*^floxed^ mice are smaller than wild type mice and exhibit cerebellar hypoplasia ^13, 18, 54^. *Cask*^floxed^;*Calb2*-Cre F1 mice were genotyped by PCR.

*Cask*^floxed^;*Calb2*-Cre mice remain indistinguishable from the *Cask*^floxed^ mice well into adulthood (∼40 days), indicating that acute deletion of *Cask* does not have significant effects on cerebellar development, motor learning, or locomotor function. Past two months of age, however, *Cask*^floxed^;*Calb2*-Cre mice begin displaying obvious locomotor incoordination and ataxia which are rapidly progressive. By approximately P100, these mice are profoundly ataxic, are unable to keep their balance and repeatedly fall over with an inability to walk forward (supplemental video). Despite profound motor coordination deficits, the *Cask*^floxed^;*Calb2*-Cre mice are otherwise healthy and display a slick coat, good body condition score, and are bright, alert and responsive. Compared to littermate *Cask*^floxed^ controls, the cerebellum of the *Cask*^floxed^;*Calb2*-Cre mouse is extremely diminished in volume at P100 when the motor phenotype has plateaued (Figure 5A, B). Comparing the histology of the *Cask*^floxed^;*Calb2*-Cre cerebellum at P30 (well before onset of ataxia) and P100 (after onset of ataxia), our results indicate that at P30, the cerebellum of *Cask*^floxed^;*Calb2*-Cre mice is populated with well-placed granule and Purkinje cells. At P100, however, we observe profound loss of granule cells, whereas Purkinje cells remain visible as a standard single layer of cells (Figure 5C). The molecular layer of the cerebellum is thin and collapsed, most likely due to loss of parallel fibers arising from the granular cells and loss of synaptic connections between granule cells and Purkinje cells (Figure 5D-G). We therefore next quantified synaptic connections within the cerebellar layers using bassoon as a pre-synaptic marker. As seen (Figure 5H-K), our data indicate that synapse density is unaltered, although the absolute number of synapses is reduced due to the shrunken volume of the molecular layer. The large number of remaining synapses are likely to be derived from the climbing fibers. Notably, other regions with Calb2-Cre recombination such as the olfactory bulb, hippocampus and striatum do not show the striking hypoplasia observable in the cerebellum. In our previous studies, we did not observe degeneration of retinal ganglion cells, which are also positive for *Calb2*-Cre ^58^. Our data here thus indicate that loss of CASK results in the disproportionate degeneration of a specific vulnerable neuronal population, cerebellar granule cells, leading to cerebellar hypoplasia.

**Figure 5.**
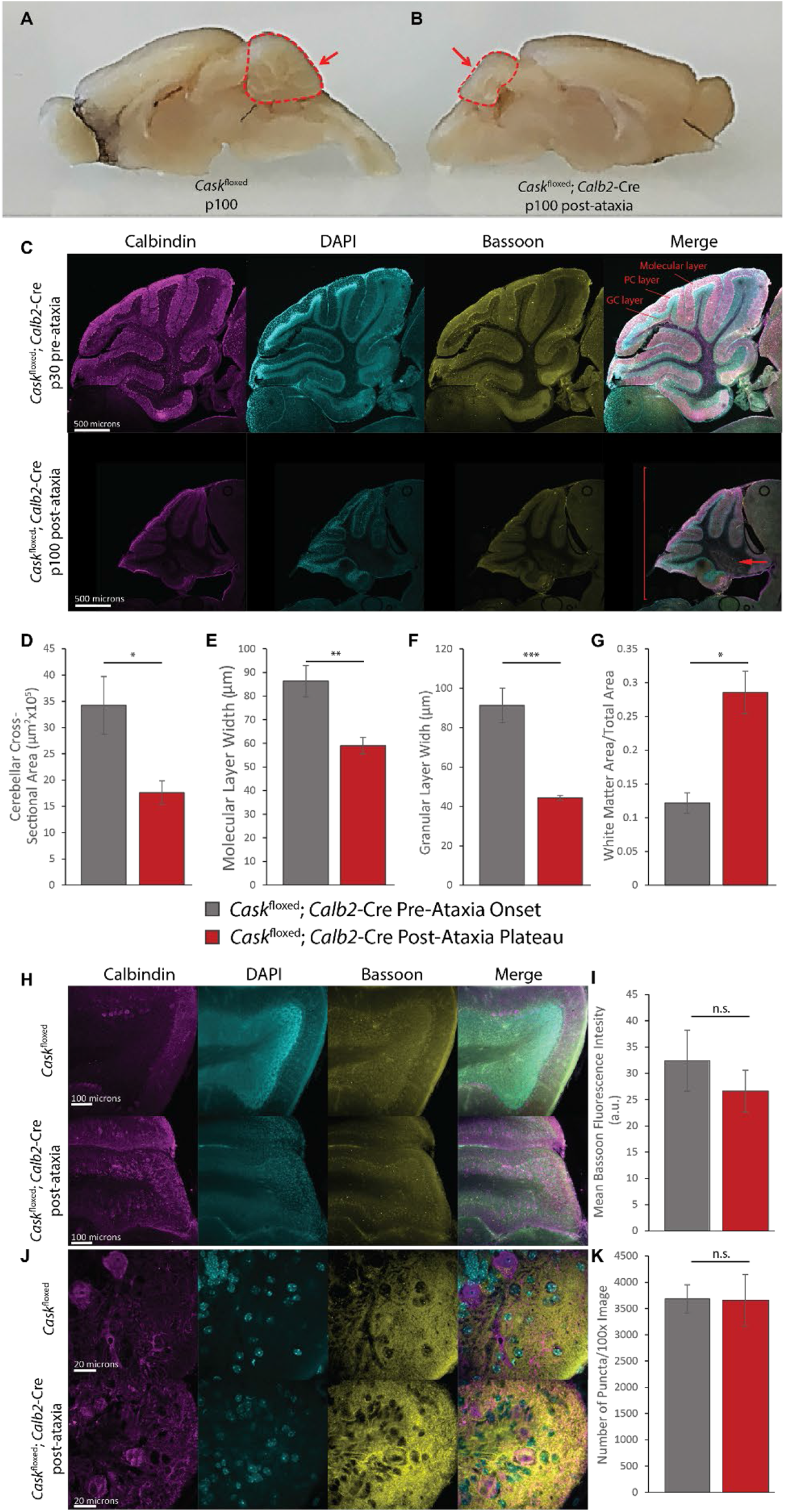
Deletion of *Cask* from post-migratory cerebellar cells results in profound cerebellar degeneration and accompanying ataxia. (A) Gross images of age-matched *Cask*^floxed^ control and (B) *Cask*^floxed^;*Calb2-*Cre brains after plateau of ataxia; arrow indicates diminished volume of the cerebellum while the remainder of the brain remains similarly sized. (C) 10x image of *Cask*^floxed^;*Calb2-*Cre at P30 (top) and P100 (bottom) demonstrating severely reduced cross-sectional area, molecular layer width and granular layer width at P100; the arrow indicates expanded white matter; the bracket indicates diminished overall cerebellar size. (D-F) Quantification of the entire cross-sectional area pre- and post-ataxia, molecular layer width, and granular layer width. (G) Ratio of white matter area to total cross-sectional area pre- and post-ataxia. n=3 for (B-G). (H) Representative 20x images of the anterior cerebellar folia in *Cask*^floxed^;*Calb2-*Cre post-ataxia plateau (bottom) and age-matched *Cask*^floxed^ controls (top) immunostained for calbindin to label Purkinje cells, DAPI to label nuclei, and bassoon to label synapses. (I) Quantification of bassoon staining intensity indicates no difference in fluorescence intensity between groups. (J) Representative 100x images of *Cask*^floxed^;*Calb2-*Cre post-ataxia plateau (bottom) and age-matched *Cask*^floxed^ controls (top). (K) Quantification of the number of bassoon positive puncta per 100x image quantified automatically using SynQuant ^80^ indicating no difference between groups. N=3 *Cask*^floxed^ and 4 *Cask*^floxed^;*Calb2-*Cre for (H-K); bars in all panels indicate mean±SEM. * indicates p < 0.05 using a two-tailed Student’s t-test.

A decrease in grey matter creates an impression of increased white matter area. On the other hand, the histopathology in the human cerebellum displayed disorganized white matter (Figure 3E). We therefore quantified myelin in the *Cask*^floxed^;*Calb2*-Cre mice using FluoroMyelin^TM^ staining. As seen in Figure 6A, the myelin appears to be disorganized in the white matter of folia from the *Cask*^floxed^;*Calb2*-Cre mouse cerebellum, which is most obvious in the region immediately distal to Purkinje cells. We also observed extremely limited myelinated axons in the anterior-most folium (Figure 6A). Quantification of pixels displayed a strong trend towards a decrease in myelinated fibers which did not reach statistical significance (Figure 6B). The degeneration of cerebellar grey matter thus is also associated with disorganization of the white matter in the *Cask*^floxed^;*Calb2*-Cre mouse cerebellum, and the broadened white matter layer is likely to be filled only with acellular matrix. Because the *Cask*^floxed^;*Calb2*-Cre mouse represents a targeted deletion of CASK in cerebellar neuronal cells, it is reasonable to conclude that the observed disordered white matter is a property of the underlying neuronal pathology rather than an oligodendrocyte-mediated pathology and confirms our observations from the human subject.

**Figure 6.**
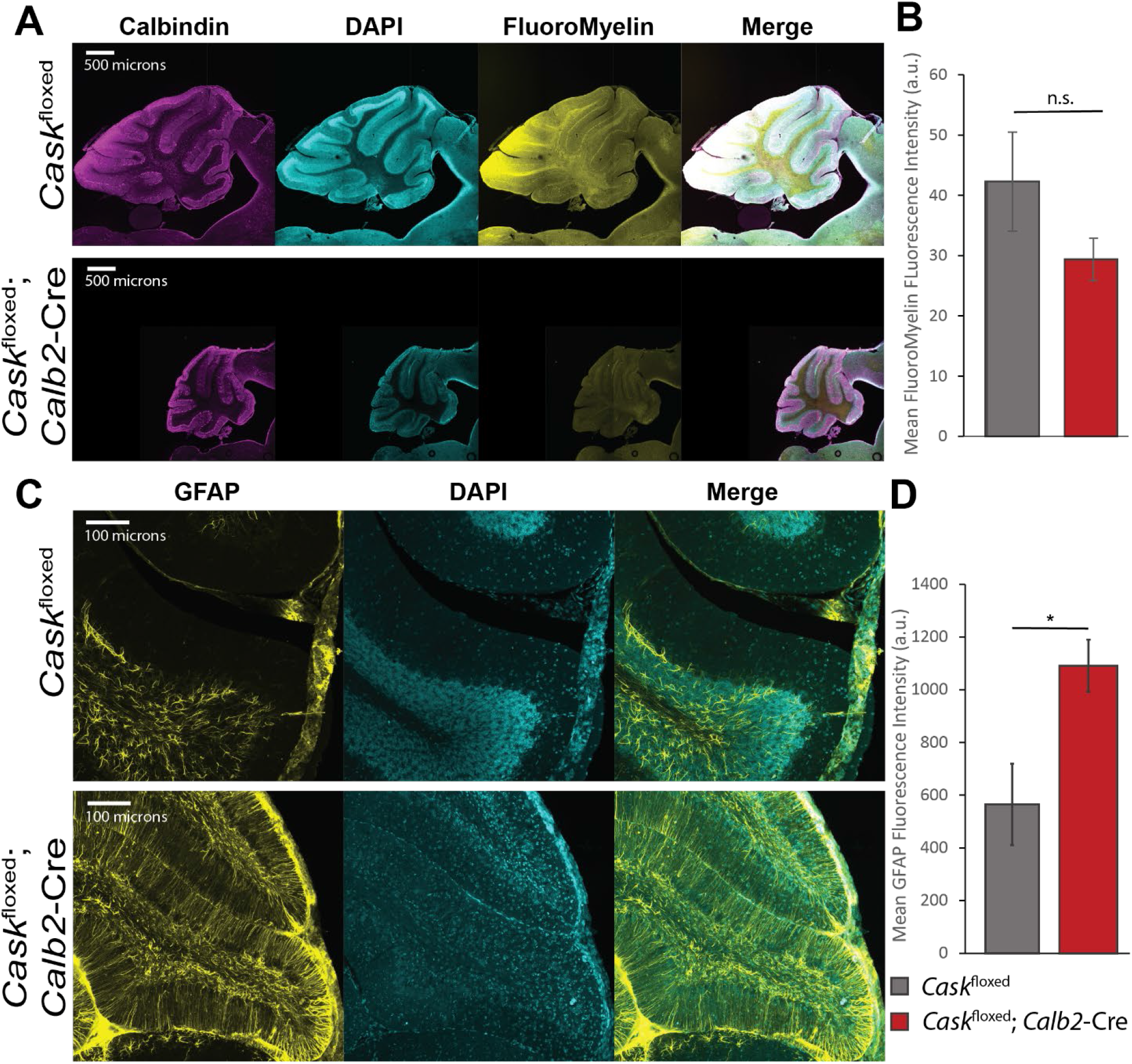
*Cask*^floxed^;*Calb2-*Cre mice display disorganized white matter and astrogliosis in cerebellum. (A) Cerebella of adult *Cask*^floxed^;*Calb2-*Cre post-ataxia plateau and *Cask*^floxed^ controls were labeled for calbindin, DAPI and FluoroMyelin (white matter). The mice display disorganized white matter with loss of myelinated fibers in the anterior folia. (B) Quantification of pixel intensity of FluoroMyelin performed with n=4 mice of each genotype. (C) Representative images of GFAP immunostaining of anterior cerebellar folia in *Cask*^floxed^;*Calb2-*Cre post-ataxia plateau and age-matched *Cask*^floxed^ controls; from left to right: GFAP, DAPI, merge. (D) Quantification of fluorescence intensity of GFAP staining by genotype; asterisk indicates p<0.05, n=3 mice of each genotype.

Neuronal loss or damage is typically associated with gliosis, as seen in the human subject, so we next immunostained mouse cerebella with a marker for reactive gliosis, glial acidic fibrillary protein (GFAP) (Figure 6C). Although *Cask*^floxed^ mice display some GFAP positivity, *Cask*^floxed^;*Calb2*-Cre mice displayed an almost 2-fold higher level of astrogliosis compared to age-matched control *Cask*^floxed^ mice (Figure 6D). Overall, this finding suggests that CASK loss-of-function produces protracted neuronal loss in the cerebellum, explaining why MICPCH typically becomes obvious a few months after birth. The cerebellar hypoplasia associated with loss of CASK represents disproportionate neuronal loss in the cerebellum.

Finally, we examined the functional loss associated with the cerebellar degeneration in *Cask*^floxed^;Calb2-Cre mice. By P100, the mouse’s hindlimbs can no longer maintain normal righting, and the mice display hindlimb clasping with no obvious dystonic movement (Figure 7A). Accelerating rotarod balance experiments suggest that even at P48, the mutant mice have a trend to underperform on a rotarod, indicating that the process of cerebellar degeneration and consequent functional degradation may be ongoing even before obvious locomotor defects are visually noticed within the cage. At P70 the mice are unable to perform on the rotarod at all, demonstrating a rapid degradation of locomotor coordination within a short span of 3 weeks (Figure 7B). Additionally, the cerebellar degeneration and accompanying motor phenotype only manifest in the homozygous knockout of CASK from cerebellar cells and not in the heterozygous deletion (Figure 7C-E) indicating that despite CASK being absent from approximately half the cells in the heterozygous deletion due to its X-linked nature, cerebellar degeneration requires total deletion of CASK. Overall, our data indicate that deletion of CASK does not affect brain development and the brain phenotype is unlikely due to defects of reelin function. Further, CASK loss leads to degeneration of cerebellar neurons leading to pronounced cerebellar atrophy that results in a progressive cerebellar ataxia.

**Figure 7.**
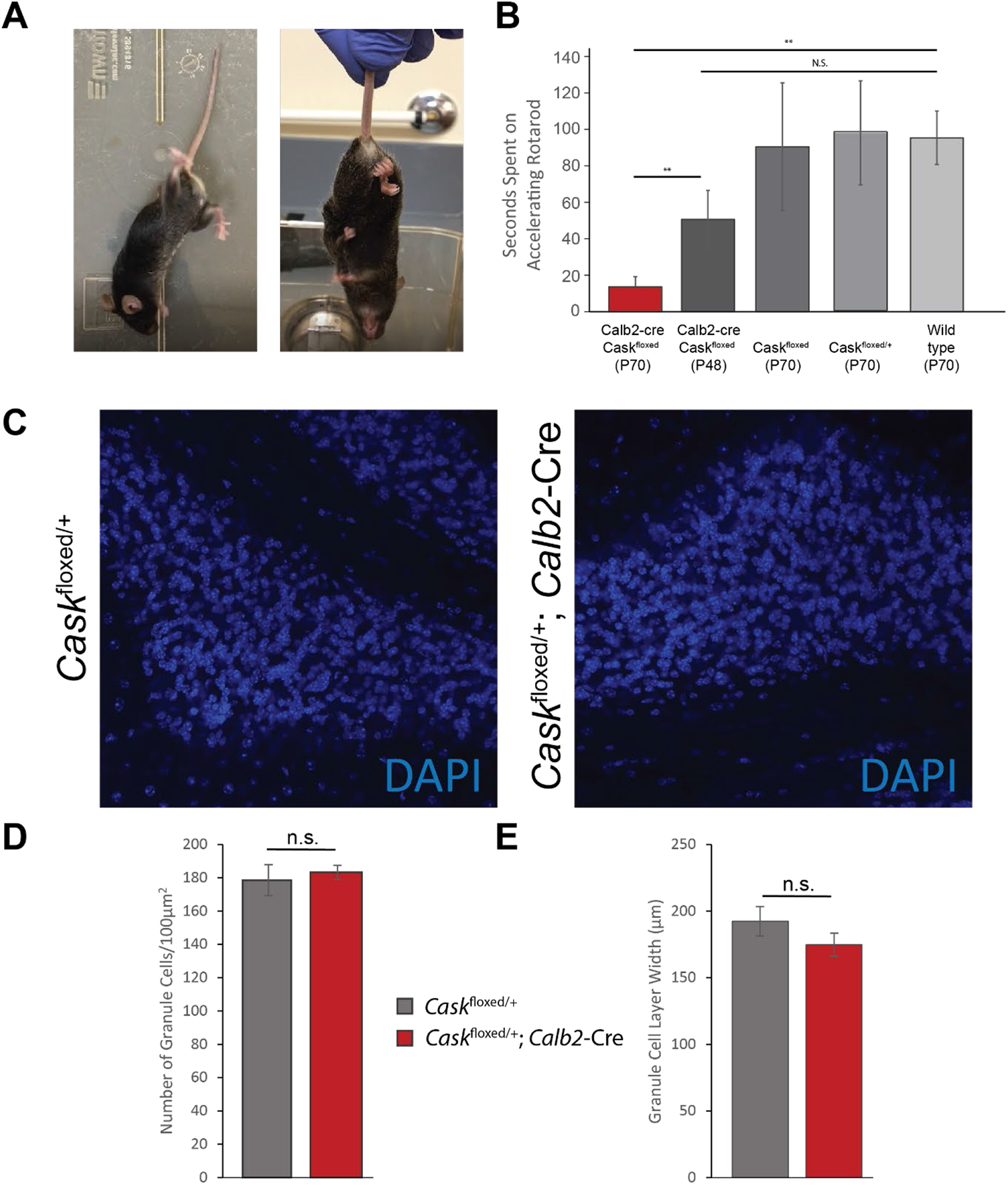
Cerebellar degeneration is accompanied by locomotor incoordination but does not occur in the heterozygous condition. (A) Example of aberrant locomotor behavior observed in a *Cask*^floxed^;*Calb2-*Cre mouse after plateau of ataxia. Example of hindlimb-clasping behavior in a *Cask*^floxed^;*Calb2-*Cre mouse. (B) Time spent on an accelerating rotarod in seconds by genotype from left to right: *Cask*^floxed^;*Calb2-*Cre post-ataxia onset (n=4); *Cask*^floxed^;*Calb2-*Cre pre-ataxia onset (n=5); age-matched *Cask*^floxed^ controls (n=4); age-matched heterozygous *Cask*^floxed/+^ controls (n=3); and age-matched wild-type controls (n=4). * indicates p < 0.05 using a two-tailed Student’s t-test. Results are plotted as mean±SEM for all panels. (C) Fluorescent images of nuclei in the granule cell layer for *Cask*^floxed/+^ control mice (left) and *Cask*^floxed/+^; *Calb2*-Cre heterozygous cerebellar knockout mice (right) aged over 1 year. (D-E) Quantification of DAPI+ nuclei density (D) and granular layer width (E) demonstrating no degenerative cell death or thinning of the granular layer compared to control in the heterozygous knockout; n=4 mice in each genotype.

## Discussion

Developmental disorders are defined based on their clinical course rather than cellular pathology. Presentation of a severe, chronic disability (mental and/or physical impairment) in three or more areas of major life activity by the age of 22 that is likely to continue through the individual’s lifetime is classified as a developmental disorder (*Developmental Disabilities Assistance and Bill of Rights Act of 2000*). Strategies for molecular therapeutic intervention, however, are more likely to be dependent on cellular pathology rather than the clinical course of a given disorder. Conditions associated with mutations in the human *CASK* gene have been described as developmental disorders ^10, 12^. *CASK* is a ubiquitously expressed gene and has been proposed to have a function in a variety of organs including the intestine, kidney, heart and brain ^21, 22^, and mutations in *CASK* produce microcephaly as well as somatic growth retardation ^10, 12^. In boys who do not express *CASK,* a clear picture has emerged consisting of neurodevastation, microcephaly, pontocerebellar hypoplasia (PCH), and a consistently abnormal EEG pattern characterized by disorganization, low frequency, attenuation and discontinuity. *CASK* null boys are thus likely to be diagnosed with epileptic encephalopathies such as Ohtahara syndrome and West syndrome. Despite uniform neurological findings in these subjects, findings involving other organ systems remain inconsistent and often unremarkable. Our analysis of *CASK* null mutations in boys indicates that the function/s of CASK that are critical for survival are brain-specific; all other organ systems can function within the normal range without CASK ^16^. In fact, we have previously demonstrated that deletion of *CASK* in neurons is sufficient to produce somatic and brain size reduction in mice ^54^. The thinning and dysfunction of the brain stem manifests as aberrant respiratory, deglutition and cardiovascular reflexes, and it is this dysfunction that underlies the lethality associated with CASK loss in mammals ^16, 18^.

Neurodevelopment could be stalled at several steps of brain development. This includes cell proliferation, neuronal differentiation and polarization, neuronal migration and final neuronal maturation including axonal and dendritic growth and synaptogenesis. At a molecular level CASK has been proposed to play a role in all of these processes. Evidence exists that suggests CASK participates in mitotic spindle orientation and cell proliferation ^61^, in cell polarization ^24^, in axonal and dendritic maturation and synaptogenesis ^40, 41, 43^. Within the synapse CASK can be found in molecular complexes that include other important molecules like Mint1, Caskin, liprin-α and the adhesion molecule neurexin ^21, 36, 62, 63^. CASK is a kinase that phosphorylates neurexin and is likely to regulate this complex formation ^62, 64–66^. The most accepted notion of CASK in neurodevelopment has, however, been the role of CASK in neuronal migration via its interaction with CINAP and Tbr-1 resulting in the upregulation of reelin transcription ^13, 45, 46, 67^. Our analysis of the CASK-null human brain is, however, unable to support any of these putative roles of CASK in neurodevelopment.

Genetic manipulation of *Cask* in murine models has demonstrated that CASK-linked murine brain pathology is postnatal and not likely to be a developmental defect ^54, 58^. Although the delivery of the decedent described here was late preterm, Apgar scores were normal, and the infant was released from the hospital without concern. Rapid regression within days to weeks has been noted in this and other boys with *CASK*-null mutations, indicating that although degeneration may begin in the third trimester, it continues rapidly throughout infancy. Despite a smaller size, the gross and histological findings in a brain without CASK are minimal. Overall, the brain configuration, vasculature, ventricular system, meninges, brain lamination and neuronal migration all remained unaltered. This indicates that in the presence of *CASK* mutation, embryonic brain development appears unchanged, with no defect in neuronal differentiation and migration. The findings from neuroimaging and histological studies of human cases are consistent with the findings of CASK knockout mice where the brain size, lamination and synapse formation are all normal at birth ^18^. In fact, absence of an acute locomotor effect in *Cask*^floxed^;*Calb2*-Cre mice excludes a synaptic role of CASK in cerebellar function. Overall our study here indicates that CASK is not likely to work in neurodevelopment via the purported Tbr-1-reelin pathway. This interpretation is in line with the observation that in the mouse model, abrogation of the CASK-Tbr-1 interaction does not affect brain size or lamination ^68^. A number of CASK missense mutations have been identified that are associated with intellectual disability and MICPCH. Recent studies on these missense mutations have also indicated that the CASK-Tbr-1 interaction is an unlikely mechanism of MICPCH ^11, 74^.

In girls with heterozygous *CASK* mutations, the predominant manifestations are also brain-related ^10, 12^. Our data presented here demonstrate that although *Cask*^+/-^ mice have cerebellar hypoplasia, they do not significantly degenerate further into old age, which agrees with the clinical definition of a neurodevelopmental disease. In girls with MICPCH and in *Cask*^+/-^ mice, however, ∼50% of cells still express CASK ^54, 67^, confounding the study of neuropathology and making it difficult to draw firm conclusions. Conditional genetic animal model experimentation allows us to overcome the difficulties presented by this X-linked condition and also helps separate the neurodegenerative pathology of CASK loss from the developmental phase.

CASK deletion in mice is lethal, but by generating otherwise healthy mice that do not express *Cask* in many parts of the brain including the cerebellum, we are able to clearly demonstrate that lack of CASK produces cerebellar atrophy and degeneration. CASK has previously been identified as a biomarker for several neurodegenerative disorders ^69–72^; our study thus explains these previous unbiased findings. CASK loss, however, does not uniformly produce neurodegeneration; for example, loss of CASK is also associated with optic nerve hypoplasia, but unexpectedly, CASK deletion from retinal ganglion cells (whose axons form the optic nerve) does not negatively affect optic nerve pathology in mice of the same genotype ^58^. Similarly, *Calb2* is present in many cortical interneurons and neurons of the olfactory bulb, hippocampus and striatum, but we do not observe death of these cells in the *Cask*^floxed^;*Calb2*-Cre mice. In fact, a closer look within the cerebellum suggests that the cerebellar degeneration may be primarily due to granule cell death, making this condition most similar to Norman-type cerebellar atrophy (OMIM: 213200). Many Purkinje cells, which are among the first cerebellar cells to express *Calb2*-Cre, do not die off but rather persist throughout the lifespan, even following the appearance of ataxia. Thus CASK loss may affect neurons differentially.

PCH pathologies are typically thought to be neurodegenerative in their origin beginning antenatally ^53^. Defects in both energy production and protein metabolism (specifically, protein synthesis) are known to disproportionately affect the cerebellum and are likely causes of PCH ^73^. In the case of CASK mutation, PCH has been described as neurodevelopmental, mostly due to the non-progressive course seen in females ^74^. In contrast to hemizygous and homozygous *Cask*^floxed^;*Calb2*-Cre mice, the heterozygous *Cask*^floxed^;*Calb2*-Cre mice fail to exhibit cerebellar degeneration (Figure 7C-E). Our findings here thus suggest that CASK-linked PCH is also neurodegenerative and the arrest of neurodegeneration in girls most likely arises from mosaicism of the defective X-linked gene, which guarantees that ∼50% of brain cells retain a normally functioning *CASK* gene (Figure 8). Mechanistically, our findings demonstrating that CASK is also likely to function in pathways associated with energy production and protein metabolism ^54, 75^, confirming that MICPCH shares not only the pathogenic mechanism, but also a common molecular pathway with other types of PCH.

**Figure 8.**
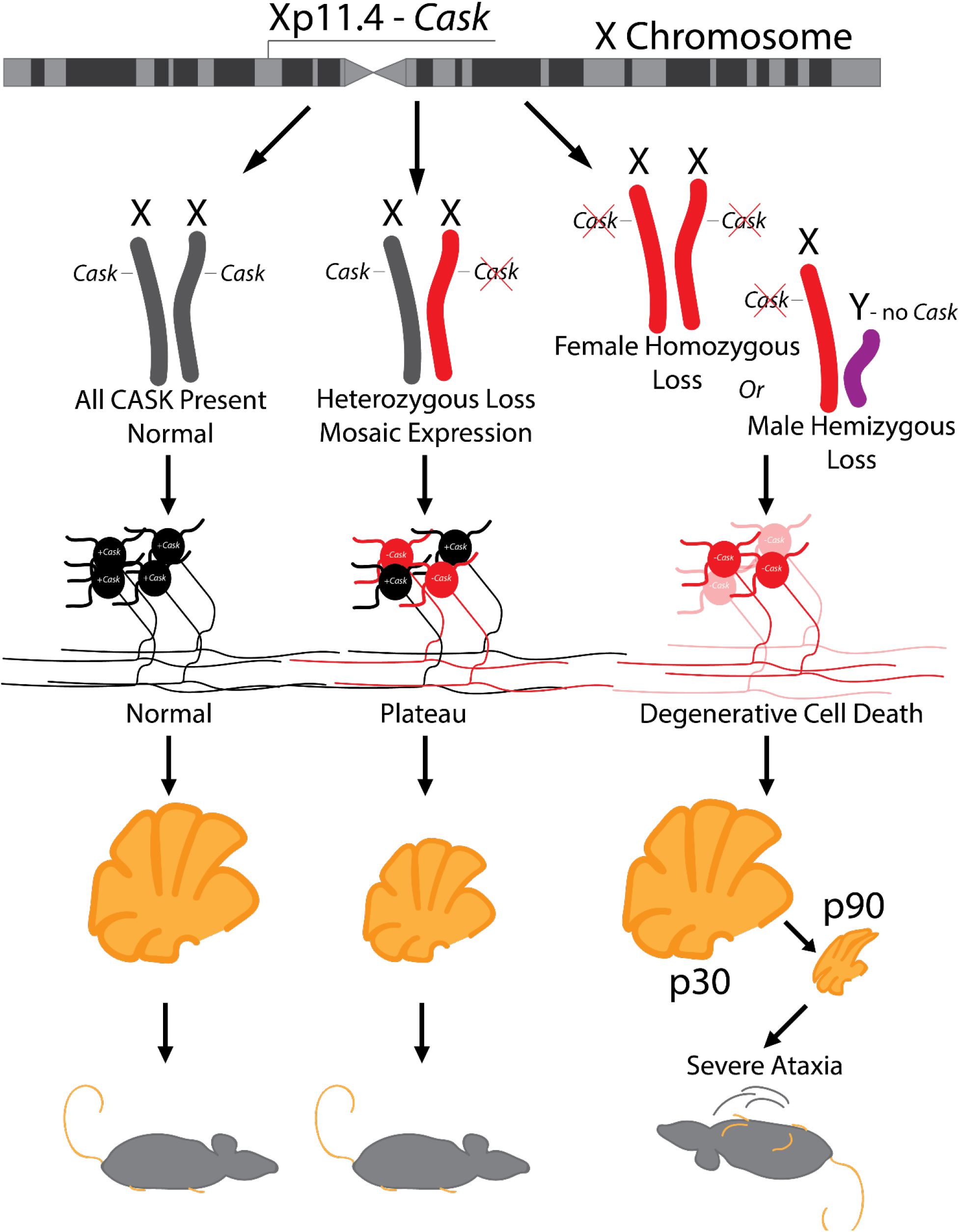
Model describing zygosity-based mechanism of a neurodevelopmental versus neurodegenerative clinical course of CASK-linked phenotype based on random X-chromosome inactivation. CASK is an X-linked gene critical for maintenance of cerebellar neurons. Heterozygous mutation in CASK produces CASK loss-of-function in only 50% of neurons (red). In the heterozygous condition (red and gray), neurodegeneration thus plateaus (bottom middle), causing an apparent neurodevelopmental disorder, whereas hemizygous CASK mutations (red and purple) in male mice or homozygous CASK mutation (two reds) in female mice produce a progressive phenotype typical of neurodegeneration with severe ataxia (bottom right).

Heterozygous mutations in X-linked genes other than *CASK*, such as MeCP2 and CDKL5, are associated with postnatal microcephaly in girls ^1^. Intriguingly, subjects with mutations in these genes show an initial normal developmental trajectory followed by developmental arrest and delay (regression) ^76, 77^. Mutational analysis of orthologous genes in murine models also produces phenotypes which are clearly post-developmental, presenting only in adulthood ^78^. Incidentally, just like CASK mutations, CDKL5 and MeCP2 mutations in males are associated with epileptic encephalopathy ^2, 3^. These data raise the possibility that even in disorders such as Rett syndrome (MeCP2 mutation) and CDKL5 deficiency, the pathology may be neurodegenerative, as originally suggested by Andreas Rett ^5^, but the clinical course may not be progressive in females due to the mosaic expression of the normal gene under heterozygous conditions.

## Materials and Methods

### Statement of ethics

All studies described herein were approved by the Virginia Tech Institutional Animal Care and Use Committee and Institutional Review Board.

### Statistics

A two-tailed Student’s t-test was used as a comparison between two genotypes in each experiment to compute significance with an alpha of 0.05.

### Clinical History

The decedent was a male born via vaginal delivery at 36.1 weeks of gestation to a 34-year-old woman, G3, P3, A1 (Gravida, para, abortus). He was conceived through in vitro fertilization with a sperm donor. He was born as a monochorionic, diamniotic twin. Ultrasound at the third trimester indicated the presence of slight microcephaly and smaller cerebellum which raised some concerns. He was also small for gestational age with a weight of 2.4 kg (0% Percentile, Z score – 7.37), length of 43 cm (4^th^ Percentile, Z score -1.71) and head circumference of 30 cm (4^th^ percentile, Z score-1.79). The Apgar scores at 1 and 5 minutes after birth were recorded as 8 and 9. The decedent was discharged from the hospital 2 days after birth. At home he became apneic with hypoventilation and was readmitted to a hospital 4 days later. Despite positive airway pressure ventilation, the apneic spells continued which led to neurological and genetic investigations. He was then diagnosed with microcephaly and pontocerebellar hypoplasia with CASK mutation. He displayed poor feeding, profound hypotonia, microcephaly, micrognathia, bilateral clubfoot, and vertical chordee with penile torsion. Oral-pharyngeal motility studies revealed mild to moderate oral motor dysphagia; there were episodes of silent aspirations with very limited reflux. A gastrostomy tube placement was performed. Fluctuations in body temperature with hypothermia and heart rate were also noted. Within 3 weeks after his birth, torso flexions were noted occurring 2-3 times a day. He also displayed tics in the hands, feet and neck which lasted for several seconds to several minutes. The decedent developed irritability and intolerance to feeds, hypothermia and acute respiratory failure with apnea. A surface, 25-channel video electroencephalography (vEEG) was performed using an international 10-20 system. A diagnosis of Ohtahara syndrome was established due to the presence of a typical burst suppression pattern. He was started on keppra and a ketogenic diet. Possibility of long-term palliative care including tracheostomy was discussed, but a decision was made against aggressive continued therapy. He passed away 2 months and 6 days after birth.

### EEG Spectral Analysis

Raw data were trimmed for artifacts by a trained observer in the clinic. Data were analyzed in MATLAB 2017a using the EEGLab toolbox. After filtering from 0.01-50Hz, bad channels were removed based on spectral power. Spectral power was plotted for each channel independently using the spectopo() function with a window length of 256 samples, FFT length of 256, and 0 overlap in the entire 0.01-50Hz frequency band. Channels covering each of a given lobe (frontal, parietal, temporal, occipital, central) were then grouped and mean power spectral density was calculated within each biologically relevant frequency band: delta, alpha, beta, theta, and low gamma. Time-frequency plots were generated for a representative 1 minute of the recording using a divisive baseline.

### Generation of Mouse Lines

*Calb2*-Cre mice (strain 010774) was obtained from Jackson Laboratory, *Cask*^floxed^ mice (strain 006382) was a kind gift from Prof. Thomas Südhof. *Cask*^floxed^ females were bred with *Calb2*-Cre positive males to generate the F1 cross *Cask*^floxed^::*Calb2*-Cre. F1 mice were bred to Ai14-LSL-tdTomato-positive males obtained from Jackson Laboratory (strain 007914) to generate fluorescent reporter mice. F1 mice were genotyped by PCR using primers targeted at either LoxP elements, a sequence within the Cre gene, or a sequence within the tdTomato gene. All lines were from a C57BL/6J background backcrossed for at least 25 generations.

### Antibodies and Material Reagents

Bassoon monoclonal antibodies were obtained from Enzo Lifescience, GFAP monoclonal antibodies were obtained from Invitrogen, calbindin polyclonal antibody from Invitrogen, synaptophysin antibody from Sigma and secondary antibodies conjugated with AlexaFluor 488, 550 and 633 were obtained from Thermofisher. Hardset Vectashield^TM^ with DAPI was obtained from Vector Laboratories.

### Immunostaining of Mouse Tissue

For all immunostaining, mice were sacrificed by trans-cardiac perfusion first with phosphate buffered saline (PBS) for exsanguination and subsequently with 4% paraformaldehyde for fixation of tissues. Brains were dissected and post-fixed for at least 24 hours in 4% paraformaldehyde. After post-fixation, brains were hemisected along the longitudinal fissure and 50µm sagittal sections were cut using a ThermoScientific™ Microm HM650V Vibratome. Sections were submerged in permeabilization/blocking solution composed of 10% fetal bovine serum and 1% Triton-X 100 in PBS overnight at +4°C.

Rabbit anti-calbindin was diluted at 1:50 in blocking solution and mouse anti-bassoon was diluted at 1:200 in blocking solution. For GFAP immunostaining, mouse anti-GFAP was diluted at 1:200 in blocking solution. After blocking/permeabilization overnight, free-floating sections were incubated for 3 hours in dilute primary antibody at room temperature. After incubation in primary antibody, sections were washed 3 times for 5 minutes in PBS before being incubated in secondary antibody for the respective host species for 1 hour at room temperature. Sections were again washed and mounted on slides using VECTASHIELD® anti-fade medium. Quantification of synapse density was conducted using the SynQuant algorithm ^80^.

### Immunostaining of Human Tissue

Human tissue obtained during autopsy was post-fixed in 10% formalin overnight, embedded in paraffin, and subsequently sectioned into 20µm sections onto charged slides. Slides were deparaffinized with 3 changes of poly-xylenes for 10 minutes each time and rehydrated using an ethanol gradient from 100%-95%-70%-50%-H_2_O for 5 minutes in each condition. Antigen retrieval was conducted by boiling slides in 10mM sodium citrate buffer with 0.1% Tween-20 for 10 minutes in a domestic microwave followed by running the slides under cold tap water for 10 minutes. Immunostaining for GFAP, calbindin and synaptophysin was then conducted using the same procedure described for mouse tissue.

### Motor Behavioral Assays

Accelerating Rotarod experiments were conducted by placing 4 *Cask*^(floxed)^::*Calb2*-Cre mice at P100 (post-ataxia onset), at P48 (pre-ataxia onset), and age-matched *Cask*^(floxed)^ control mice on an accelerating Rotarod, beginning at 2 cycles/minute and accelerating at a rate of 5 cycles/minute until mice fell off the platform. Three trials were conducted in succession for each mouse, with 5 minutes of rest between trials.

## Author contribution

Experiments were conceived by KM, PP, MF, IC. Data analyzed by, KM, MF, PP, JH, LL, SS. Experiments conducted by PP, AK, JH. Papers written by KM, PP, MF, LL, SS, IC, JH.

## Acknowledgements

The work was supported with funding from the NIH National Eye Institute (R01EY024712) to KM. We thank Drs. Thomas Südhof and Alexei Morozov for providing CASK*^floxed^* mice and Ai14-LSL-tdTomato mice, respectively.

## Notes

### Competing Interest Statement

The authors have declared no competing interest.

