## Supplemental Figures for "Complete loss of CASK causes severe ataxia through cerebellar degeneration in human and mouse"

**Supplemental File:**

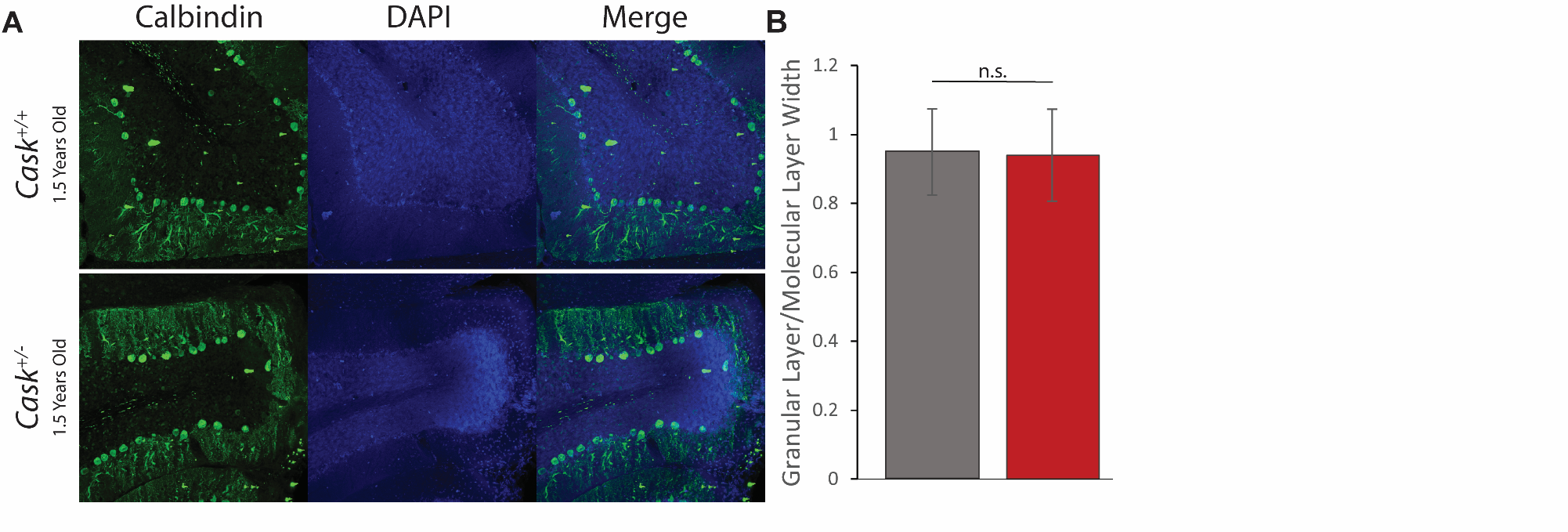
Supplemental Figure 1. (A) Representative images of cerebella from *Cask*^+/-^ mice (bottom) and *Cask*^+/+^ littermate control mice (top) aged up to 1.5 years. (B) Quantification of the ratio of the width of the granular layer over the molecular layer demonstrating no diminishment of granular layer width in the heterozygous absence of CASK even at extremely advanced ages. N=3 mice for each genotype.

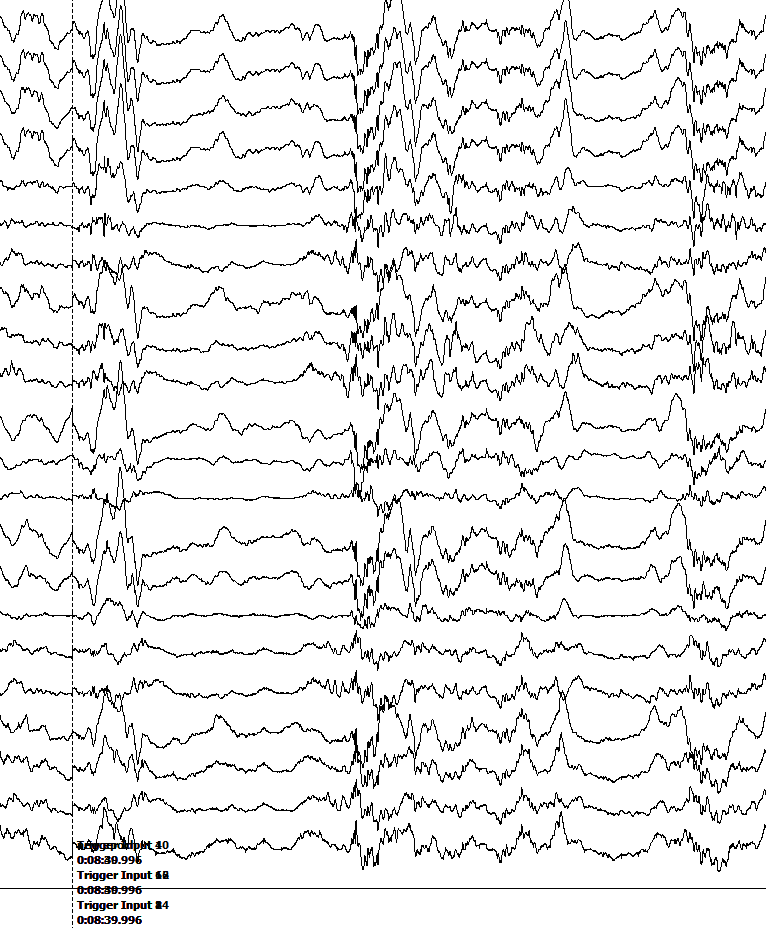

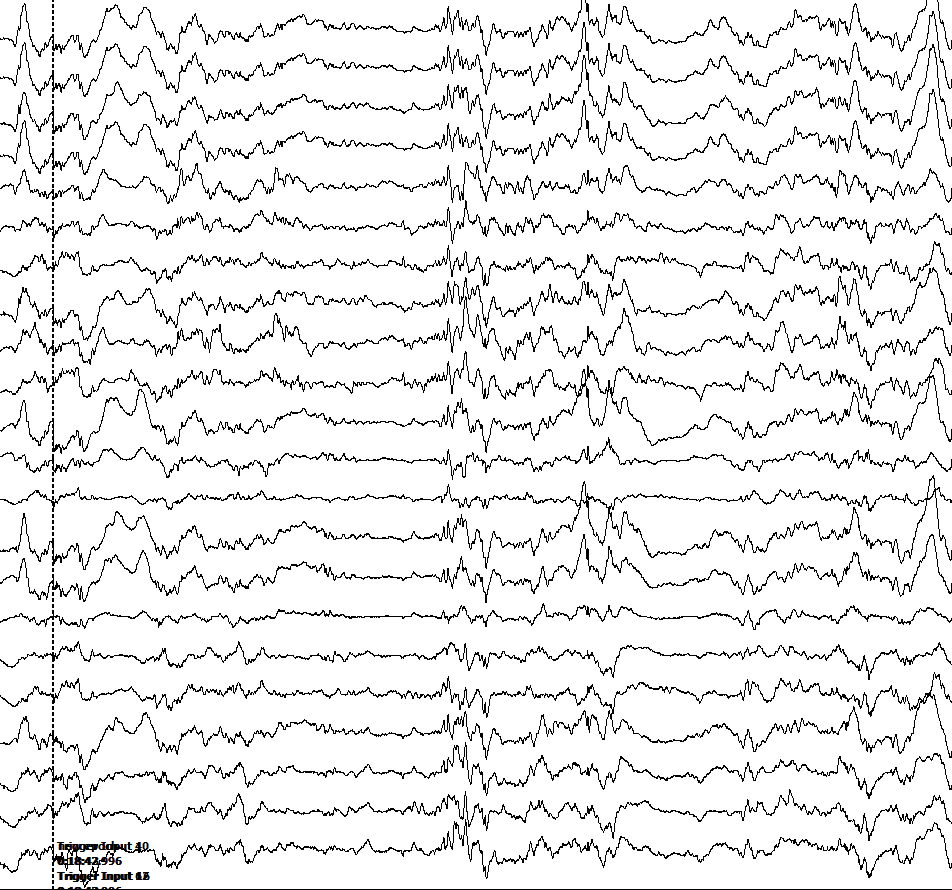

Supplemental Figure 2. Examples of burst-suppression pattern in EEG recordings.

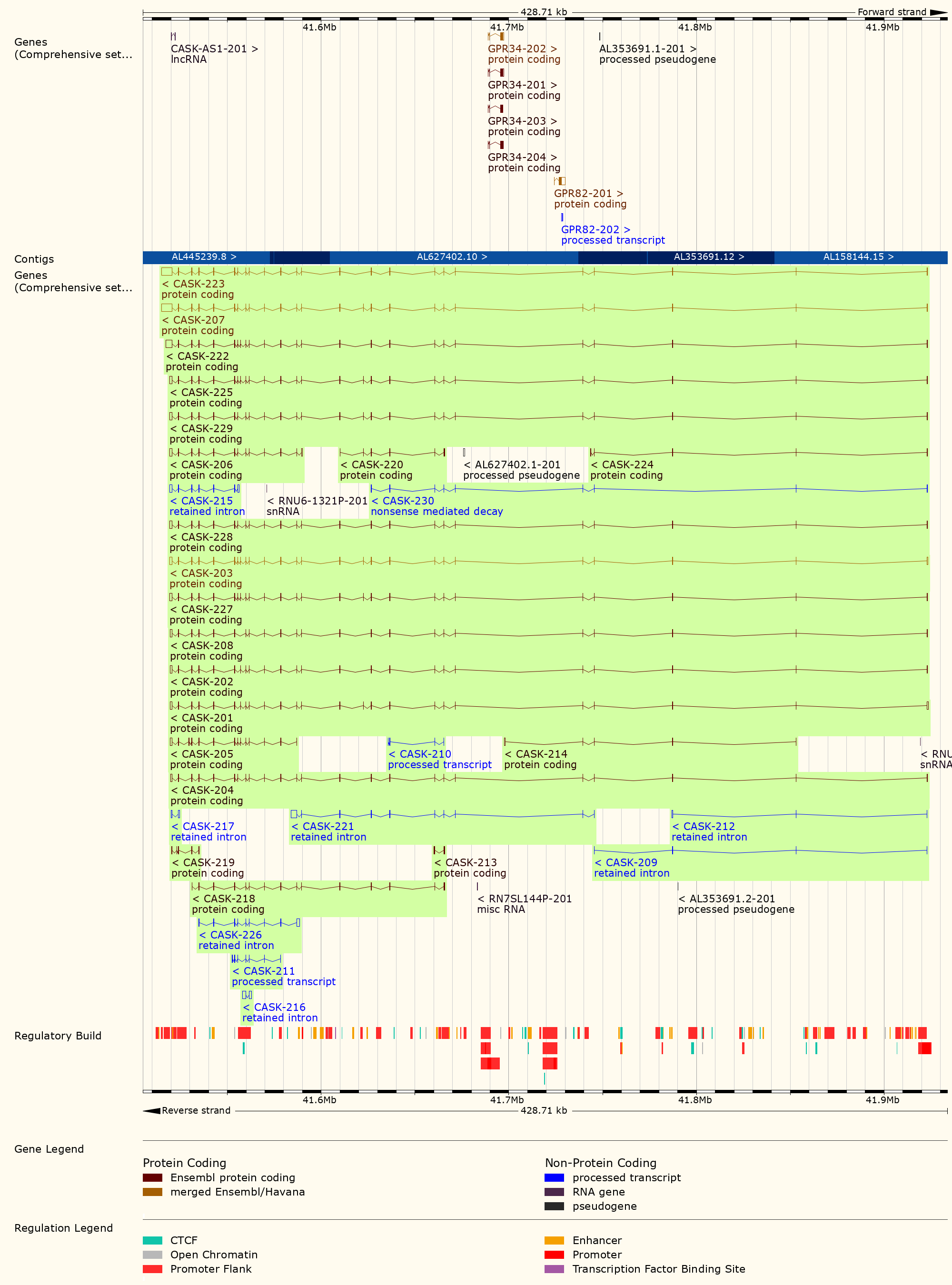

Supplemental Figure 3. NCBI summary of human *CASK* gene (top) and known splice variants (bottom). All functional transcripts include the second exon.

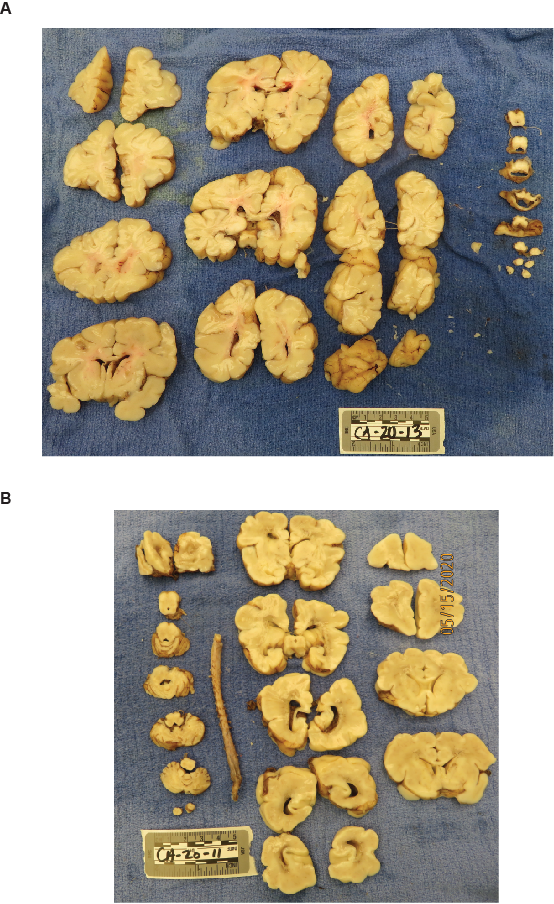

Supplemental Figure 4. Macroscopic view of 1 inch coronal slices of brain from (A) *CASK* R27Ter subject (2 months) (B) a child who died of non-neurological cause (40 days old). Note the disproportionate cerebellar and brain stem hypoplasia. The grey and white matter configuration remains normal in *CASK* R27Ter subject.

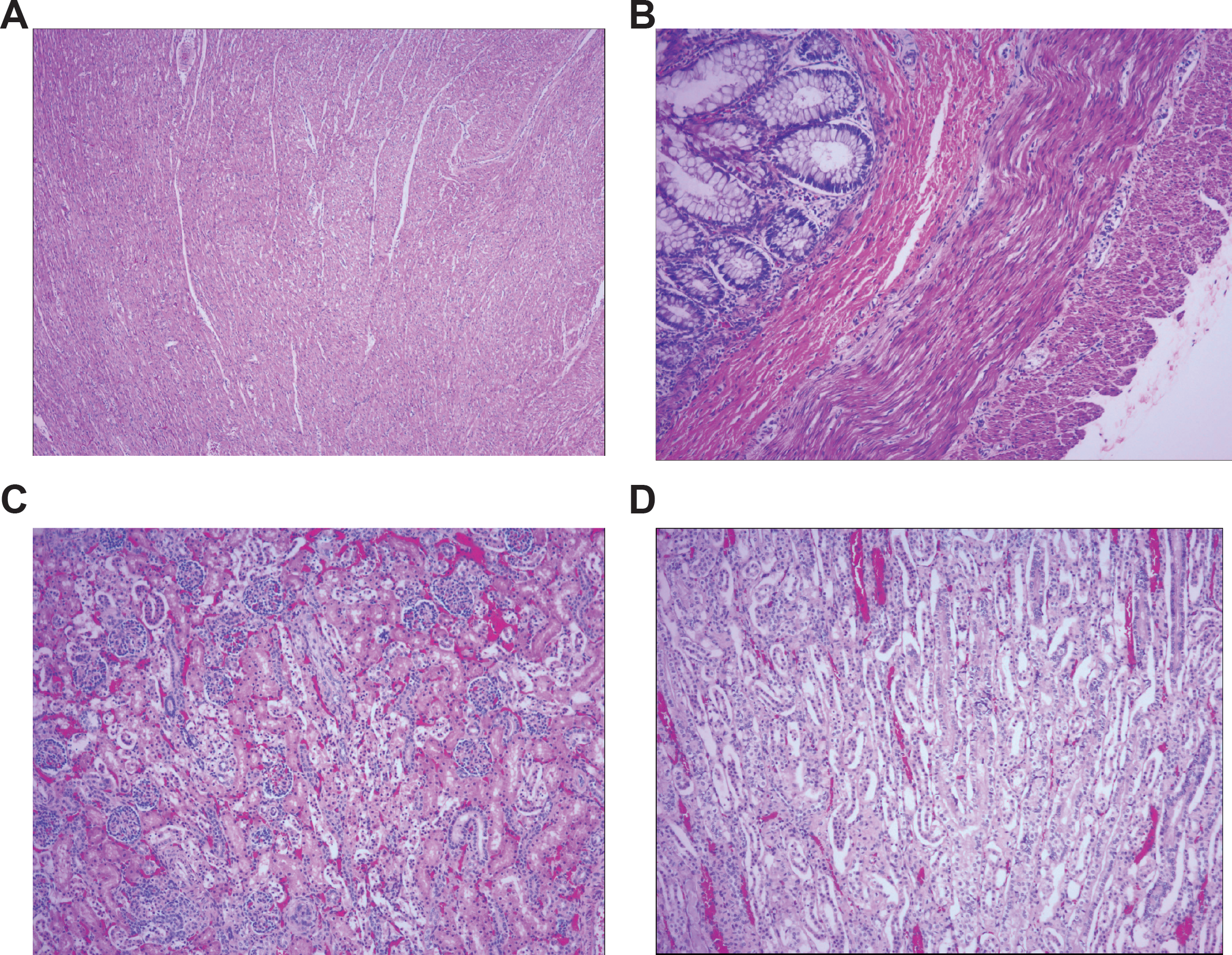

Supplemental Figure 5. Hematoxylin and eosin stain of (A) heart showing uniformly layered myofibers with no obvious pathology (4X), (B) rectum showing mucosal, submucosal, muscularis externa and serosal layers, neuronal plexuses are visible with no obvious pathology (10X), (C) renal cortex and (D) renal medulla showing normal glomeruli formation and renal tubules (10X).

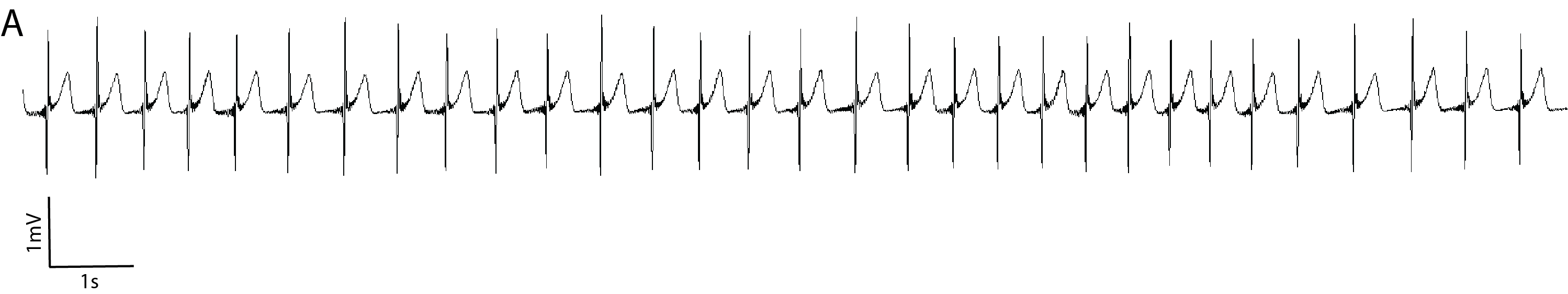

Supplemental Figure 6. (A) Representative 20 second electrocardiogram trace (precordial chest lead) demonstrating a sinus rhythm.
